# Physical properties and actin organization in embryonic stem cells depend on differentiation stage

**DOI:** 10.1101/2020.04.27.063891

**Authors:** K. G. Hvid, Y. F. Barooji, I. Isturiz, J. M. Brickman, L.B. Oddershede, P. M. Bendix

## Abstract

The cellular cytoskeleton provides the cell with mechanical rigidity and mediates mechanical interaction between cells and with the extracellular environment. The actin structure plays a key role in regulating cellular behaviors like motility, cell sorting, or cell polarity. From the earliest stages of development, in naïve stem cells, the critical mechanical role of the actin structure is becoming recognized as a vital cue for correct segregation and lineage control of cells and as a regulatory structure that controls several transcription factors. The ultrastructure of the earliest embryonic stem cells has not been investigated in living cells despite the fact that it is well-known that cells undergo morphological shape changes during the earliest stages of development. Here, we provide 3D investigations of the actin cytoskeleton of naïve mouse embryonic stem cells (ESCs) in clusters of sizes relevant for early stage development using super resolution optical reconstruction microscopy (STORM). We quantitatively describe the morphological, cytoskeletal and mechanical changes appearing between cells in small clusters at the earliest stages of inner cell mass differentiation, as recapitulated by cells cultured under two media conditions, 2i and Serum/LIF, thus promoting the naïve and first primed state, respectively. High resolution images of living stem cells showed that the peripheral actin structure undergoes a dramatic change between the two media conditions. The actin organization changed from being predominantly oriented parallel to the cell surface in 2i medium to a more radial orientation in Serum/LIF. Finally, using an optical trapping based technique, we detected micro-rheological differences in the cell periphery between the cells cultured in these two media, with results correlating well with the observed nano-architecture of the ESCs in the two different differentiation stages. These results pave the way for linking physical properties and cytoskeletal architecture to the development from naïve stem cells to specialized cells.

**Statement of Significance:** Cells receive mechanical signals and must provide mechanical feedback, therefore, physical properties are instrumental for cell-cell interactions. Mechanical signals mediated through the cell surface can significantly affect transport of signaling molecules and can influence biological processes like transcriptional regulation. To achieve a deeper insight into how the cytoskeletal structure is responsible for cell shape and material properties at the earliest stages of development, we employ super-resolution microscopy to image actin fibers in clusters of embryonic stem cells mimicking early development. By modification of the culturing conditions, we investigate how the actin cytoskeleton and micro-rheological properties of ESCs change between the naïve ground state and the stage primed towards epiblast, thus revealing a correlation between differentiation stage and cytoskeletal structure.

## INTRODUCTION

The biomechanical properties of cells allow them to sense and react towards their physical environment, for example by sensing the rigidity of the environment. The mechanical properties, both of the cell and its environment, play an important role in cell organization, migration and differentiation. During development, the physical properties of cells change drastically in terms of viscoelasticity and morphology (1), with the cell generally becoming softer, more compliant, as it progresses in development. Furthermore, the rigidity of the environment has been shown to be crucial in differentiation, for regulation of stem-ness of cells, and for the development of organoids (2). The viscoelasticity of cells is also important in disease, for example in cancer where invasiveness has been linked to the cells’ stiffness and adaptability to extracellular matrix stiffness (3, 4). As the exact mechanisms and effects of both mechanical properties and mechano-sensing of cells during differentiation and development are not yet well understood, it is important to map and understand the changes in physical properties of cells during differentiation and to correlate this with mapping of the differential expression of transcription factors. Optimally, we need to understand the interplay between mechanical and biochemical cues during differentiation.

The changes in the overall viscoelasticity of cells undergoing differentiation has been investigated (1, 5-7) and cells are known to become mechanically progressively stiffer the closer they are to pluripotency, thus indicating differences in their cytoskeletal organization (8). However, little information exists on the local sub-cellular viscoelastic properties of ESCs and the associated changes in the cytoskeletal structure as they differentiate. The actin cortex in ESCs has been imaged and shown to be regulated via a complex interplay between Arp2/3, formin, and capping proteins while myosin II was proven to be only minimally involved (5).

Embryonic stem cells are known to exhibit dramatic changes in morphology and mechanics during the first few days after fertilization. Cultured ESCs can be maintained in a ground state of totipotency or pluripotency by maintaining certain culture conditions and their exact stage can be detected by associated expression levels of transcription factors (9, 10). Cells cultured in serum-free media with the addition of two small molecule inhibitors of GSK3 and MEK along with leukemia inhibitory factor (LIF) resemble the 3.5 day post fertilization (11-13). These cells exhibit a greater level of pluripotency than cells grown in serum media supplemented with LIF (9, 13, 14) and exhibit a remarkably different morphology when grown on a substrate. To our knowledge, no study has quantitatively investigated the structural and physical properties of cells cultured in these two widely used culture conditions.

Here, we investigate mouse embryonic stem cells (ESC), which recapitulate cells from the developing embryo, however, they can also be cultured and expanded *in vitro*, and studied in detail at the sub-cellular level. To address early development, we culture ESCs in two well-described stem cell media conditions; (i) a complete media based on LIF and fetal bovine serum (FBS) referred to as Serum/LIF and (ii) a media known as 2i/LIF and here referred to as 2i. Cells grown under both conditions mimic cells in the inner cell mass (ICM) of the mouse blastocyst about 3.5 days post-fertilization and just before the ICM starts differentiating into epiblast and primitive endoderm lineages (Fig. 1A,B). However, cells grown in 2i show biological (transcription and differentiability) characteristics of a more naïve stage than cells grown in Serum/LIF (15). The specific stage of each cell in the blastocyst under development is particularly interesting from a biomechanical perspective, as the differentiation into epiblast and primitive endoderm is accompanied by a physical segregation of the cell lineages into an inner and outer layer. In this study we aimed to describe whether, on the single cell level, there would be differences in either actin cytoskeletal structure or viscoelastic properties of cells grown in either 2i or Serum/LIF, respectively, representing cells in two different and early differential stages.

**FIGURE 1.**
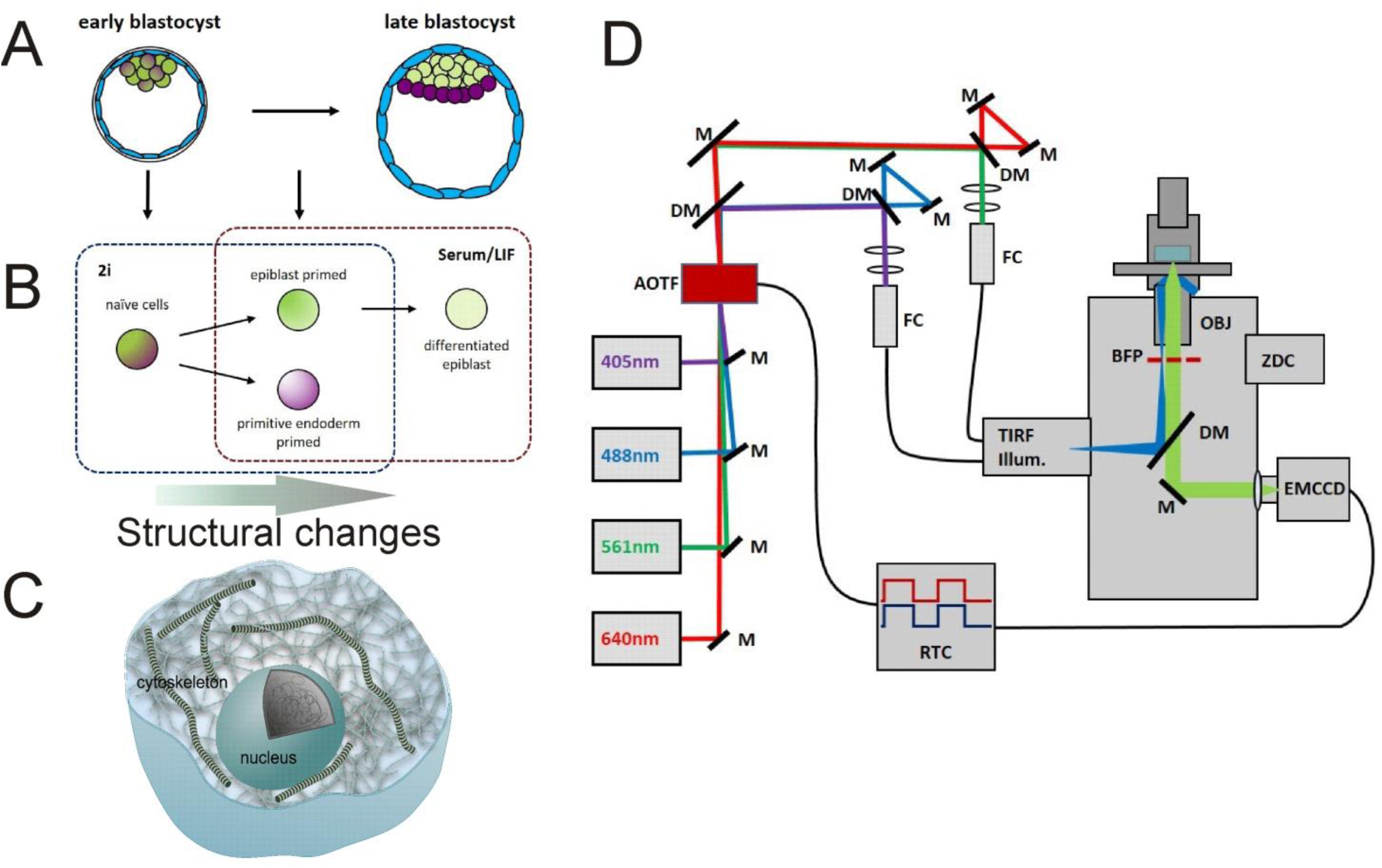
Embryonic stem cell development and the super-resolution imaging platform used to image the actin organization during early development. **(B)** Overview of how the differentiation stage of embryonic stem cells can be controlled by culturing in different media. Cells in 2i recapitulate a slightly earlier and more naïve stage of embryonic development than ESCs cultured in Serum/LIF. **(C)** Schematics showing the cytoskeletal structure of a cell. The ultrastructure of actin is not constant during development. **(D)** Description of the TIRF/STORM setup used to image actin organization in embryonic stem cells. Optical elements and light path in the TIRF/STORM setup: *M*, mirror, *DM*, dichroic mirror, *AOTF*, acousto-optical tunable filter, *FC*, fiber coupler, *TIRF Illum*., motorized TIRF illumination module, *BFP*, back focal plane, *OBJ*, objective, *ZDC*, real-time z-drift compensation module, *EMCCD*, electron-multiplying CCD camera, *RTC*, real-time controller. See Fig. S1 for more details.

To address these questions, we used both confocal imaging and super resolution fluorescence imaging of the actin structure in cells grown in 3D in small colonies. Confocal imaging was used to characterize the 3D shape of the cytoskeleton in ESC colonies grown under different culture conditions. Confocal imaging gives information on the morphology of individual cells in the 3D cluster as well as the overall distribution of actin in the cell, while it has a relatively good penetration depth in the axial direction, however, the optical resolution restricts it from being able to resolve the detailed structure of the dense network of filaments. To overcome this limitation and to examine the nanoscale structure of subcellular actin filaments, we implemented stochastic optical reconstruction microscopy (STORM) (Fig. 1D and Fig. S1), using the stochastic blinking of fluorophores linked to actin and individually detecting and localizing these molecules with nanoscale precision, thus mapping out the structure actin network (16-19).

To correlate the observed changes in cytoskeletal architecture with the micro-rheological properties of the individual cells, we probed the viscoelastic properties of live cells with sub-cellular resolution by optical trapping endogenously occurring lipid granules inside the cells. This method allows for precise quantification of the viscoelastic properties of the cytoplasm at the sub-cellular level (20).

## MATERIALS AND METHODS

### Cell culture

E14Ju mouse embryonic stem cells (ESCs) were maintained in either full stem cell media with serum and LIF (Serum/LIF) or serum-free 2i/LIF media (2i), for no more than 25 passages, and were regularly tested for mycoplasma.

Serum/LIF medium consisted of GMEM (Sigma-Aldrich) supplemented with 10% Fetal Bovine Serum (FBS), 2 mM L-Glutamine, 1 mM Sodium Pyruvate, 0.1 mM 2-mercaptoethanol, 0.1 mM Non-Essential Amino Acids, and 1000 units/mL LIF (DanStem, Copenhagen, Denmark).

2i medium consisted of a 1:1 mix of DMEM F/12 (Gibco) and Neurobasal Medium (Gibco), supplemented with N2, B27, 100 μM 2-mercaptoethanol (Sigma-Aldrich), 1000 units/mL LIF (DanStem), 3 μM Chir99021 (Sigma-Aldrich), 1 μM PD0325901 (Sigma-Aldrich).

Cells were grown on flasks (Corning) coated with 0.1% gelatin (Sigma-Aldrich) in 37 °C incubators containing 5% CO_2_, and were passaged with DPBS (Sigma-Aldrich) and Accutase (Sigma-Aldrich).

### Cell fixation and staining

For confocal and STORM imaging, cells were plated on fibronectin coated #1.5H thickness 8-well glass bottom slides (Ibidi, Gräfelfing, Germany). Fibronectin coating was performed by covering the glass slides in a solution of DPBS with 10 μg/mL human fibronectin (EMD Millipore, Burlington, MA, USA) for at least 2 hours. Cells were then plated at a density of 20,000 cells/cm^2^, and left growing in specific culturing media (as described above) overnight.

For fixation, before performing confocal and STORM microscopy, each well was washed once with DPBS before adding 4% paraformaldehyde in PBS for 10 minutes. After fixation, cells were again washed and stored in DPBS at 5 °C until staining.

Before staining, cells were permeabilized using 0.1% Triton X-100 (Sigma-Aldrich) in PBS for 15-30 minutes. After washing in PBS, blocking was performed by incubating the sample with 1% BSA in PBS for 30 minutes at room temperature. Cells were then stained with Alexa Fluor 647 Phalloidin (Invitrogen) at a 40X dilution of the stock solution for 20 minutes at room temperature. The cells were again washed in PBS before finally staining chromatin with the DAPI stain NucBlue Fixed Cell ReadyProbes Reagent (Invitrogen).

### Confocal microscopy

Fixed and stained cells were imaged on a Leica SP5 confocal microscope (Leica Microsystems, Wetzlar, Germany) using a 63X water-immersion objective (NA=1.20, Leica).

### Micro-rheology

Micro-rheological measurements were carried out using a tightly focused laser beam, an optical trap, implemented in the confocal microscope. The trapping laser had a wavelength of 1064 nm and was coupled in through the side port, the setup is described in Ref. (21). For these experiments the cells were kept alive in a perfusion chamber with the lower coverslip coated by fibronectin. As tracer particles, we used endogenously occurring lipid granule which are abundantly present throughout many cell types (20, 22, 23). The laser was focused on a single granule located at the position of interest as detected by bright field imaging. The forward-scattered laser light was collected by a high numerical aperture oil immersion condenser and imaged onto a quadrant photodiode (QPD) located close to the back focal plane of the condenser. Data from the QPD were analyzed by a custom made Matlab code based on the software published in Ref. (24) and the methods described in (23).

Briefly, the time series were Fourier transformed and for each time series the exponent, *α*, describing the scaling properties of the power spectrum in the frequency interval 300-3000 Hz was extracted. In our experiments, the laser was operated at low laser powers and this frequency interval was well below the corner frequency characterizing the stiffness of the trap, also, in this frequency interval, thermal fluctuations dominate over non-equilibrium activity within living cells (25). For anomalous diffusion, such as the thermal fluctuations of a tracer particle within a crowded cytoplasm, the mean squared displacement (MSD) of the tracer particle scales with time-lag (*τ*): *MSD*(*τ*) ∼ *τ*^*α*^, where *α* is the scaling exponent (22). For 0 < *α* < 1 the movement is sub-diffusive (26) and within this regime, the lower the value of *α*, the more elastic the environment and the closer to 1, the more viscous the environment. For pure Brownian motion *α* = 1, and 1 < *α* is a sign of super-diffusive motion. For tracer movement in a viscoelastic medium, the power spectrum scales with frequency, *f*, as:

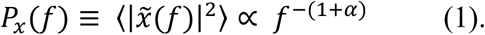

Hence, the scaling exponent, *α*, characterizing the viscoelastic properties of the medium, can be found by fitting eq. (1) to the power spectrum (20, 23) in an appropriate frequency window.

### STORM imaging and analysis

STORM imaging was based on a custom made total internal reflection (TIRF) Olympus IX83 inverted microscope (Olympus, Tokyo, Japan), equipped with a 150X 1.45 NA oil-immersion objective (Olympus), an EMCCD camera (Hamamatsu, Hamamatsu, Japan), a Z drift compensation system (IX3-ZDC2, Olympus), and a motorized TIRF module (Olympus). To enable STORM imaging, we individually fiber coupled a 640 nm laser (Toptica Photonics, Gräfelfing, Germany) and a 405 nm laser (Olympus) to the TIRF module, and independently set both illumination angles to a TIRF penetration depth of 100-200 nm for imaging. More details of the setup can be found in the Supplementary Material.

To induce stochastic blinking of the Alexa Fluor™ 647 fluorophore, imaging was performed in a PBS based imaging buffer based on the dSTORM buffer described by van de Linde and others (19) containing 100 mM MEA (Sigma-Aldrich), 0.6 mg/mL Glucose Oxidase (Sigma-Aldrich), 10% (w/v) Glucose (Sigma-Aldrich), and 60 μg/mL Catalase (Sigma-Aldrich) in PBS. The pH of the buffer was adjusted to between 7.5 and 8.5 by addition of KOH.

Time-lapse imaging of blinking events was performed at 20 ms exposure time for 10000-30000 images. The exposure time is chosen such that a sufficient signal can be collected, but without getting multiple blinking events from the same location. Individual single molecule blinking events were then localized in the time-lapse images using the ThunderSTORM (27) plugin in Fiji (28), and drift was corrected using the cross-correlation function in ThunderSTORM.

### 3D STORM imaging

3D STORM imaging was done as described above, with the important difference that a cylindrical lens was inserted in the optical pathway before the camera. This lens produced astigmatism in the detected emission from single molecule blinking events, elongating them in either the vertical or horizontal direction depending on their z-distance from the focus plane (29). The elongation was compared to the elongation from a control experiment where surface-fixed beads were scanned while gradually moving them through the focus in the z-direction. This allowed for 3D localization of blinking events using ThunderSTORM and the elongation-correlation dataset.

### Image analysis

All quantifications were done using custom written Matlab programs Matlab (Mathworks, Natick, MA).

The area and the aspect ratio (the ratio between major and minor axis) of adherent cells were calculated using Matlab and imageJ software.

The contact angles shown in Fig. 2C were calculated as the angle between the substrate (horizontal) and the edge of the cell colonies. For each colony, the angle was measured along the entire circumference of the colony and averaged. The contact area (as depicted in Fig. 3E) was measured as the total area covered by actin for individual cells detected within the TIRF volume.

**FIGURE 2.**
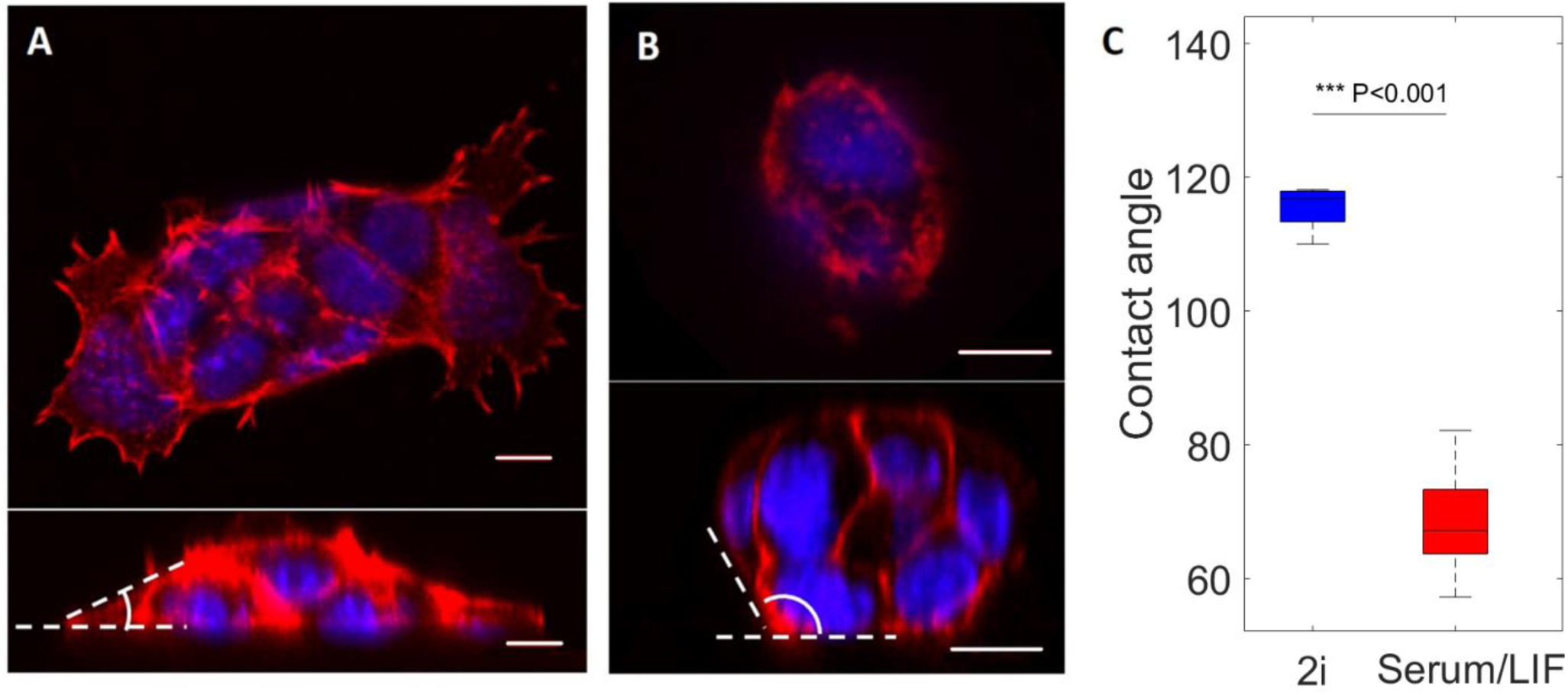
Confocal images and analysis of the shape of ESC colonies grown in 2i versus Serum/LIF. **(A**,**B)** Upper panels: images in the lateral plane (x,y). Lower panels: images from a side view (x,z) of typical cell colonies of cells grown in Serum/LIF medium (A) or 2i medium (B), respectively. The contact angle of the colony to the substrate is marked in white. The scale bar is 10 μm in all images. **(C)** Contact angle of ESC colony to the surface in 2i or in Serum/LIF, respectively, extracted from side-view images as shown in A and B. n=5 colonies for each condition, colonies grown in 2i exhibit significantly larger contact angles than those from in Serum/LIF. Error bars denote one standard deviation, and the squares show the mean values. P value was found using standard t-test.

**FIGURE 3.**
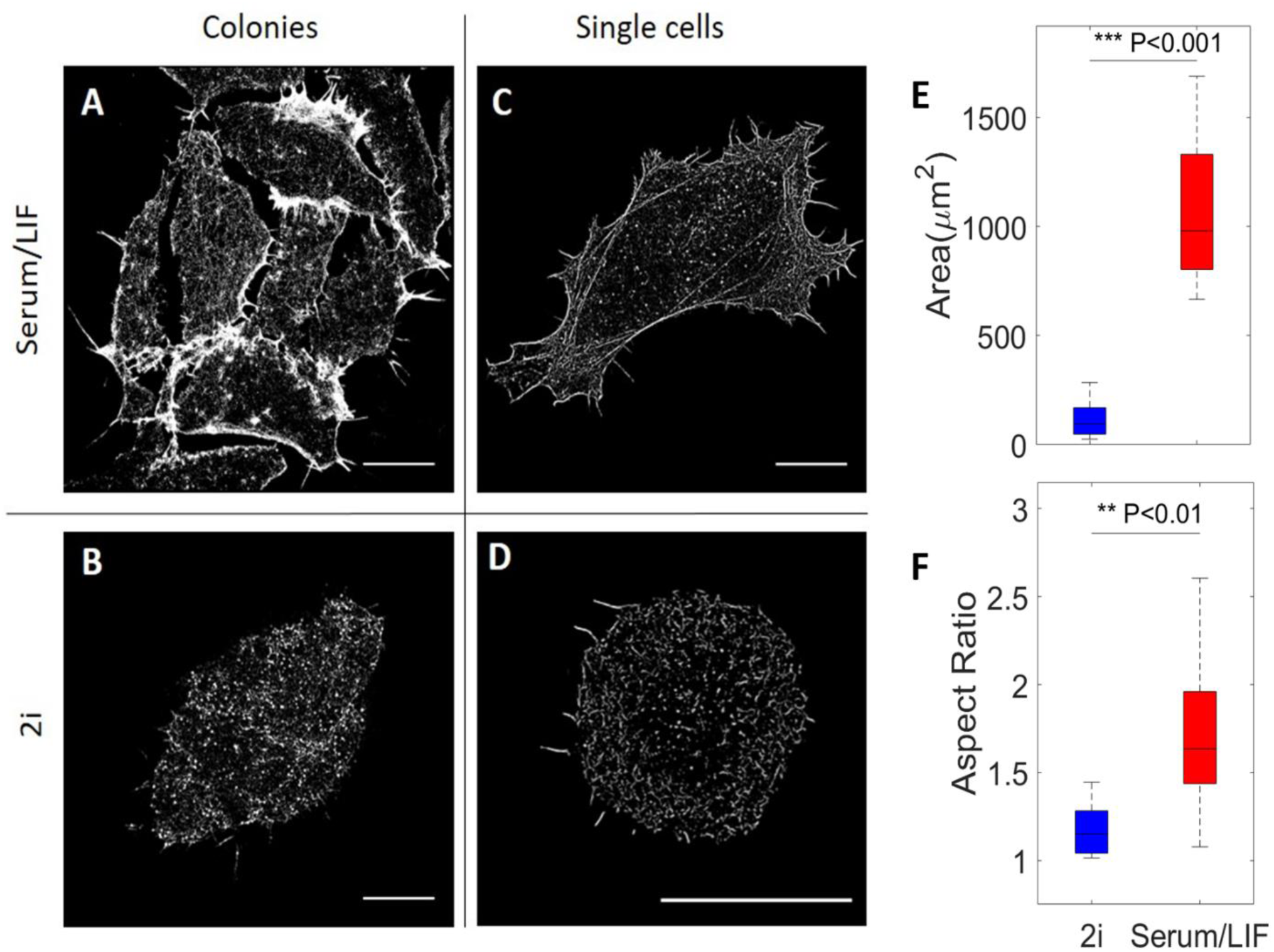
Super resolution based analysis of actin cytoskeleton organization in stem cells near the substrate. **(A-D)** STORM images of the actin network under different conditions: **(A)** a colony of cells grown in Serum/LIF media, **(B)** a colony of cells grown in 2i media, **(C)** a single cell grown in Serum/LIF media, and **(D)** a single cell grown in 2i media. All scale bars are 10 µm. **(E)** Boxplot of the surface spreading area of ESCs grown in either 2i (blue) or Serum/LIF (red) media, respectively. There is a significantly larger spreading of cells grown in Serum/LIF media than in 2i (n=16, for each condition). **(F)** Boxplot of measured aspect ratios of ESCs grown in 2i (blue) or Serum/LIF (red) media, respectively (n=16, for each condition). Cells cultured in Serum/LIF are significantly more elongated than those in 2i. Box edges indicate 25th and 75th percentiles, and whiskers extend to the most extreme data points not considered as outliers.

For quantifying actin cable orientation (as shown in Fig. 5), STORM images were filtered using an implementation of a Gabor filter to enhance the spatial resolution of linear structures in images (30, 31) (see Fig. S2). After image enhancement, the local orientation angles of actin filaments in every pixel were extracted based on the structure tensor method using the freely available ImageJ plugin OrientationJ (32, 33). Figs. 5A and 5B show the color-coded angle distribution for two cells cultured in Serum/LIF and 2i media, respectively. The polar histograms of actin orientation, from Figs. 5A and 5D, are shown in Figs. 5C and 5D. The distribution of actin orientations indicates that actin filaments are predominantly orientated along the cell’s major axis direction for a cultured cell in Serum/LIF and distributed randomly for a cell in 2i. This observation can be quantified by an orientation order parameter, *S* = ⟨cos(2*θ*)⟩, where *θ* is the local angle between the filaments in a given pixel and the direction of the cell’s major axis (34, 35). Order parameters, *S*, for the cells as depicted in Fig. 5A and B were extracted for 16 cells in each medium (Fig. 5E).

**FIGURE 4.**
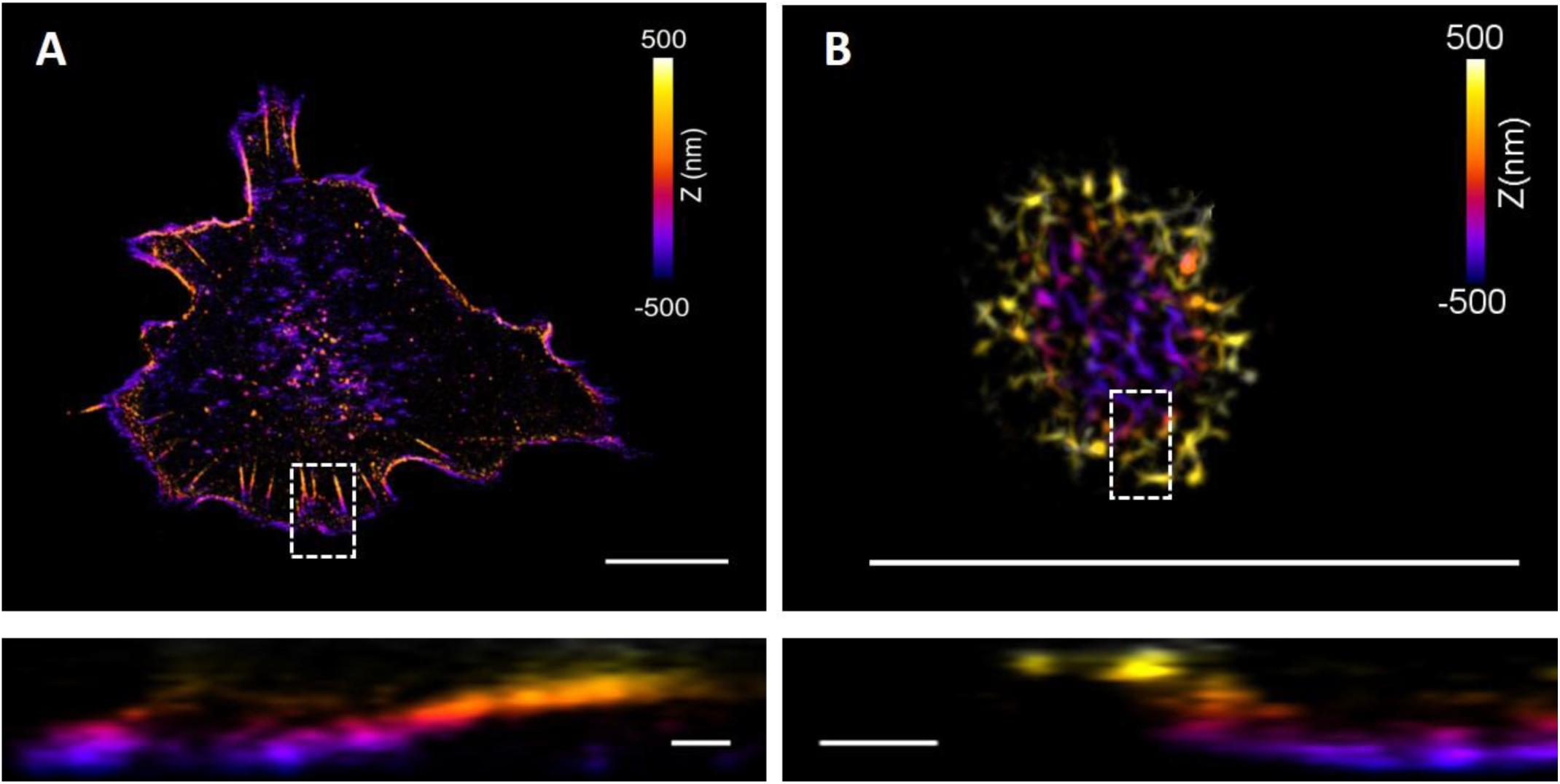
3D STORM imaging of single cells for resolving actin and stress fiber organization in ESCs grown in Serum/LIF or 2i media, respectively. **(A)** 3D visualization of actin filaments in stem cells grown in Serum/LIF. **(B)** 3D visualization of actin filaments grown in 2i. The z-positions are color-coded (violet indicating the substrate). Lower panels show side views of the boxed regions in the images above. The images indicate a dorsal type of stress fibers extending out from the surface at the edge of the cells to be present (only) in cells grown in Serum/LIF. In contrast, actin fibers in cells grown in 2i appear to extend away from the substrate close to the cell’s periphery. Scale bars in upper panels 10 μm, in lower panels 500 nm.

**FIGURE 5.**
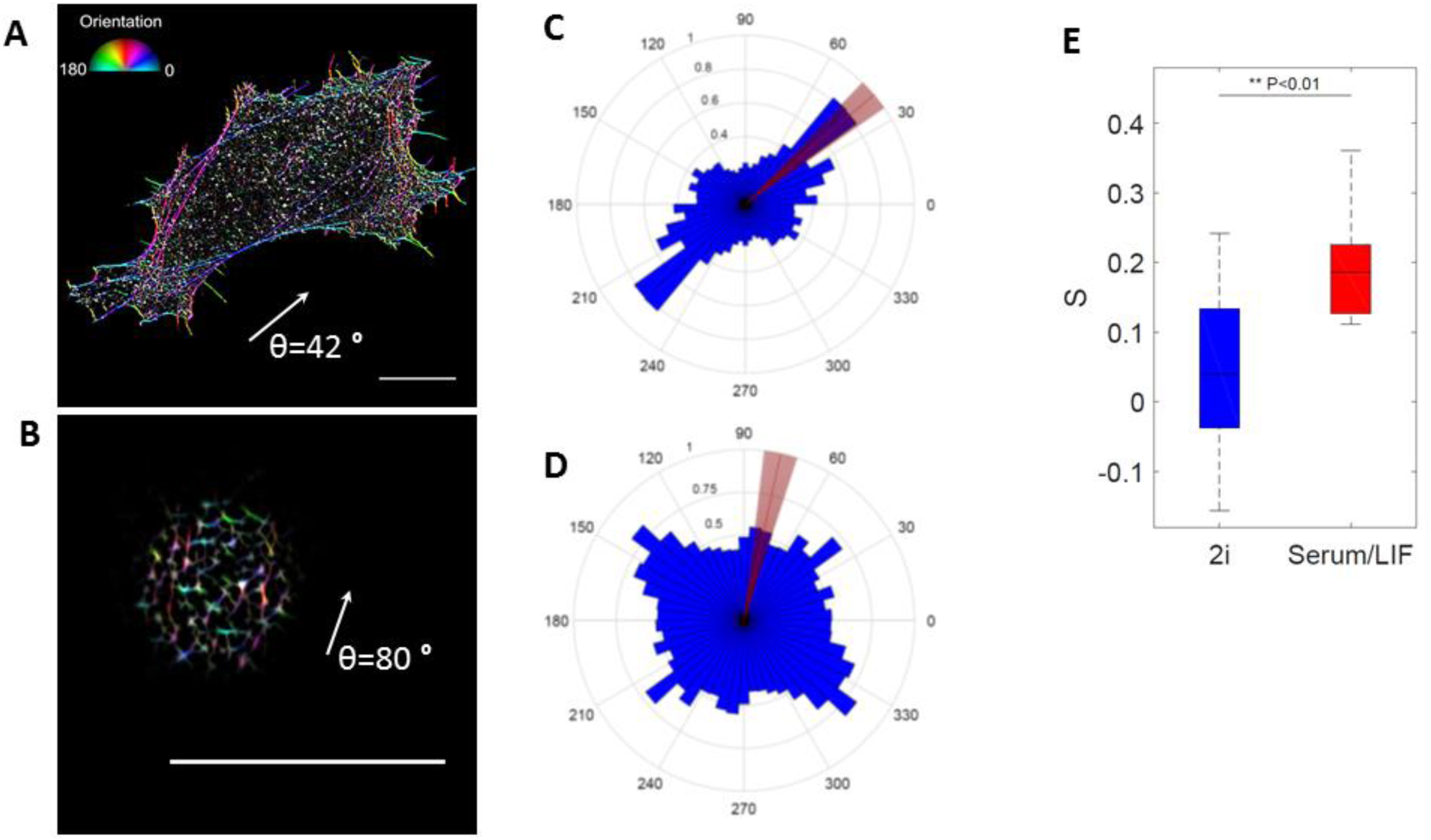
Characterization of actin filament orientation in ESCs near the surface. **(A-B)** Orientation of actin filaments in ESCs, color-code indicates orientation with respect to the cell’s major axis (white arrow), **(A)** shows an ESC grown in Serum/LIF and **(B)** an ESC grown in 2i media. Scale bars: 10 µm. **(C-D)** Angular plots showing the direction (blue) of the actin filaments with **(C)** corresponding to image **(A)**, and **(D)** corresponding to image **(B)**, respectively. The red lines indicate the orientation of the cells’ major axes found by fitting an ellipse to the adherent area. **(E)** Boxplot of the actin filament orientation order parameter *S* = ⟨cos(2*θ*)⟩, where *θ* is the angle between fiber orientation and the orientation of the cell’s major axis, Box edges indicate 25th and 75th percentiles, and whiskers extend to the most extreme data points not considered outliers. n=16 cells for each of the 2i and Serum/LIF categories. ** indicates p<0.01 using t test.

## RESULTS

### Geometry of stem cell colonies depend on stage of differentiation

To perform confocal imaging of colonies of cells in three dimensions we labeled fixed cells for F-actin and with a nuclear stain (Fig. 2A and 2B). Actin was observed to mainly localize in the cell cortex both in cells cultured in Serum/LIF and in 2i, however, with important differences: i) cells cultured in Serum/LIF have a higher number of filopodia and actin protrusions along the surface than those cultured in 2i, ii) colonies of ESCs cultured in 2i grow in rounded relatively high dome-like colonies as has previously been observed (36, 37), whereas cells cultured in Serum/LIF media grow in flat and more spreading colonies of cells, as apparent from the x-z projection (Fig. 2A,B).

At the colony edges, the contact angles of cells adhering to the substrate under the two culturing conditions were significantly different. The contact angles, defined as the angle between the substrate and colony edge and averaged over the circumference of the colony (Fig. 2C), were found to be *θ* = (115 ± 3) ° for cells grown in 2i, and *θ* = (69 ± 9) ° for cells grown in Serum/LIF. Hence, cells cultured in Serum/LIF media seem to maximize their contact with the surface by spreading out towards the substrate, whereas cells in 2i tend to minimize their contact to the substrate interface while the colony forms a dome-like structure.

### Nanoscale architecture of actin depends on stage of differentiation

To investigate the nanoscale organization of F-actin, we performed STORM imaging at depths up to a few hundred nanometers using TIRF illumination (Fig. 3). With its higher resolution than confocal imaging, STORM imaging revealed that cells cultured in Serum/LIF have concentrated actin filaments in dense stress fibers and bundles that are mainly located on and along the circumference of the cell. Actin is also clearly visible in the abundantly present filopodia. Within the central region of Serum/LIF cells, we did not observe connected filaments within the TIRF volume, indicating a lack of connected filaments close to the surface. In comparison, actin in clustered 2i cells (as shown in Fig. 3B) is more evenly distributed throughout the cell cortex at the substrate interface (within the TIRF volume).

To compare the spreading of cells in the two different differentiation stages, we quantified the contact area of each cell with the substrate. This was defined as the total area of cells visible within the TIRF volume (Fig. 3E). We found that ESCs grown in Serum/LIF media exhibit significantly larger spreading on the surface than cells grown in 2i, corresponding well with the more spread-out and flatter geometry of colonies grown in Serum/LIF than in 2i (as visible in Fig. 2). Also, as quantified in Fig. 3F, cells grown in Serum/LIF have a significantly higher aspect ratio and hence exhibit a more elongated shape than cells grown in 2i.

STORM imaging of colonies of cells grown in 2i revealed clear gaps of a few microns at the surface plane between cells (Fig. 3B). More irregular and less clear gaps are also visible between cells in a colony grown in Serum/LIF (Fig. 3A). However, colonies grown in 2i have wider gaps which appear to result from the more spherical shape of single cells and a larger contact angle formed with the surface by each of the cells cultured in 2i. These gaps could not be resolved by confocal imaging under the same conditions, indicating that actin polymers in individual cells grown in 2i are curving inward towards the center of the cell at the substrate (as indicated in the sketch in Fig. 6A). To obtain higher 3D resolution than achievable by confocal microscopy, we investigated the structures by 3D STORM (Fig. 4). For colonies grown in 2i media, we did indeed see the actin curving away from the surface at the edges of the cell. This is in contrast to the situation for cells grown in Serum/LIF, where actin bundles can clearly be seen close to the substrate and pointing towards the edges or in the protrusions of the cell (Fig. 4A). This is consistent with the presence of dorsal stress fibers in primed cells, attaching to the substrate at the leading edges of the cell (38, 39). For cells grown in 2i media, on the other hand, we do not see evidence of dorsal stress fibers, instead an interconnected mesh of actin without any clear directions or bundles is observed (Fig. 4B).

**FIGURE 6.**
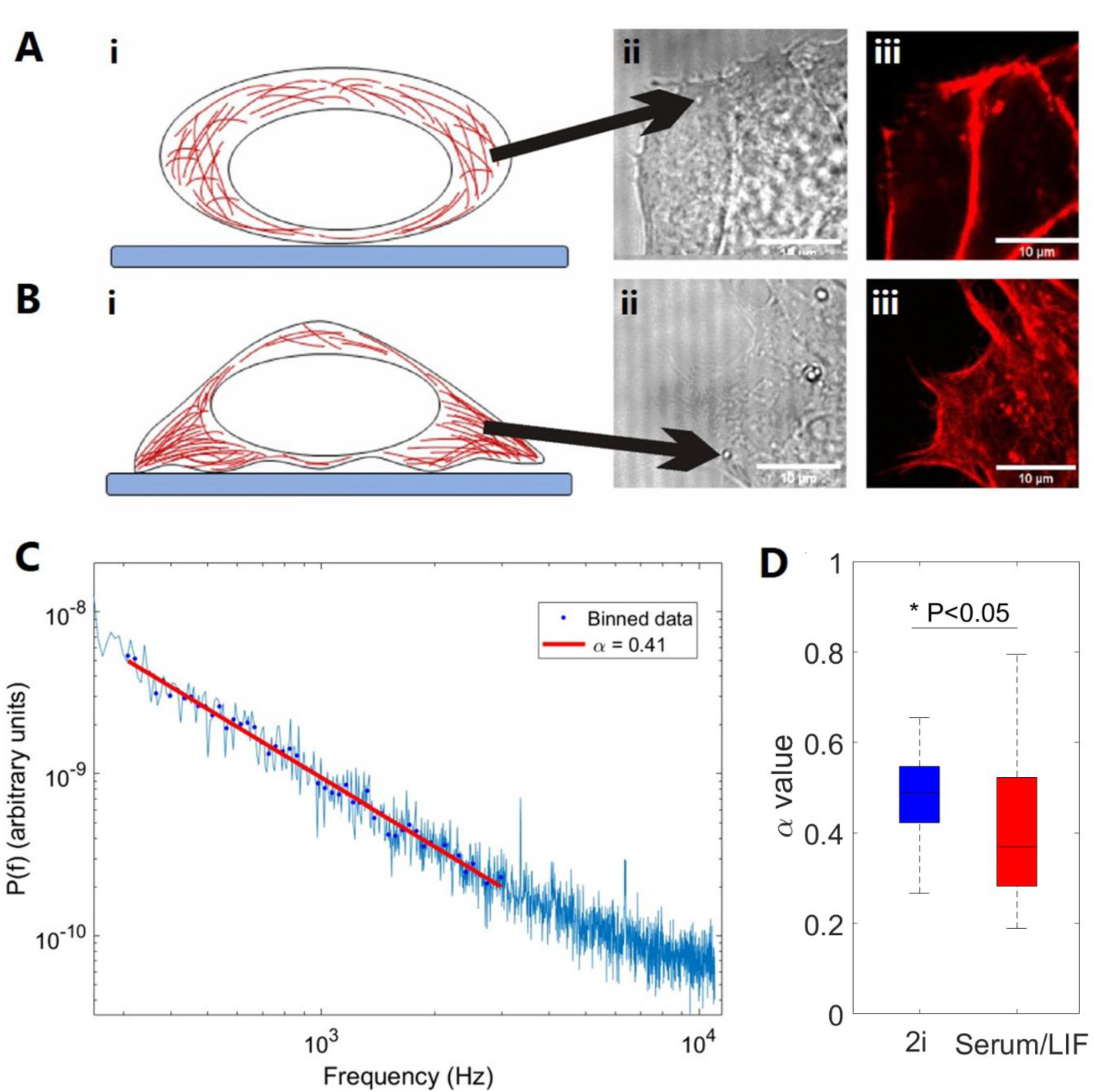
Characterization of viscoelastic properties of ESCs grown in 2i or Serum/LIF at the sub-cellular level close to the cell periphery. **(A-B)** i) Schematic of observed cell geometry, ii) bright field image, trapped granule marked by black arrow, iii) confocal images of same area as shown in ii) with F-actin labeled by a SIR-actin In **(A)** cells were grown in 2i, in **(B)** cells were grown in Serum/LIF. **(C)** Representative power spectrum of the positions visited by a lipid granule close to the periphery of an ESC cultured in Serum/LIF. The red line shows the fit of eq. (1) to data in the frequency interval 300-3000 Hz, returning α =0.41 for this experiment. **(D)** α-values from lipid granule trapping measurements in 2i or Serum/LIF, respectively. Box plot of 5th to 95th percentile, * indicates p<0.05 using t-test. N=26 cells for each condition.

### Nanoscale organization of actin at the surface depends on media conditions

In order to quantify the seemingly disorder in the orientations of filaments in 2i, we used 2D STORM data, and implemented a Gabor filter to emphasize and connect the STORM point detections into filaments. The filaments were then color coded based on their orientation respective to the major axis of the cells (Fig. 5). For each culturing condition, we quantified the orientation order parameter *S* = ⟨cos(2*θ*)⟩, where *θ* is the angle between local fiber orientation and the overall cell orientation (Fig. 5C). Actin filaments in stem cells cultured in 2i were found to be significantly more disordered than filaments in Serum/LIF cells (p<0.01). The order parameters, *S*, for the cells as depicted in Fig. 5A and 5B were found to be 0.23 and 0.05, respectively. The same analysis was done for 16 cells in each medium and the average result shows a significant difference in the order parameter: For cells cultured in Serum/LIF the average was found to be <S>=0.19 whereas for cells cultured in 2i <S>=0.04 (Fig. 5 D).

### Comparing viscoelastic properties in the periphery of cells grown under different culturing conditions

As a significant difference in actin structure of ESCs was observed under the two different culturing conditions, in particular in areas close to the cell periphery, we next set out to probe the viscoelastic properties inside the live cells. More precisely, using optical tweezers we quantified the viscoelastic properties in a frequency interval of 300-3000 Hz where non-equilibrium processes have been shown to be negligible (25) and material properties are probed at timescales relevant for the dynamics of cytoskeletal elements. In this frequency interval, the positional power spectrum, *P*_*x*_*(f)*, scales with the scaling exponent, *α*, which carries information on the viscoelastic properties as outlined in Materials and Methods.

Figure 6C shows an example of a positional power spectrum from an endogenous lipid granule located at the cellular periphery of an ESC cultured in Serum/LIF, hence, a situation resembling the one shown in Fig. 6B ii. For each culturing condition, 26 individual granules were trapped and their scaling exponent fitted and plotted in the boxplot shown in Fig. 6D. For cells cultured in 2i, we found an average scaling exponent of *α* = 0.48 ± 0.11, while for cells cultured in Serum/LIF we found an average value of *α* = 0.40 ± 0.15. In both cases only granules close to the cell periphery were investigated. First, a D’Agostino and Pearson normality test was performed on the two distributions to validate normality, second, a Student’s t-test was performed on the two distributions which returned a value of p=0.03. Hence, on a 0.05 percent significance level the two distributions would likely be independent. In conclusion, we find that close to the cellular periphery cells cultured in Serum/LIF are more elastic (stiffer) than cells cultured in 2i, which are more viscous (softer). This corresponds well to the fact that STORM imaging revealed significantly denser and straighter actin filaments to be present along the periphery of cells cultured in Serum/LIF than in cells cultured in 2i. Averaging over the entire cell, we observed no difference in viscoelastic properties between the cells cultured under the two conditions, only at the periphery a difference was observed.

## DISCUSSION

Confocal imaging of ESCs at two early developmental stages revealed clear differences in the morphology both of single cells and of the colonies they form. Cells grown in 2i, corresponding to the totipotent state, had a more spherical shape and formed dome-like cell clusters, resembling the 3.5 day cell stage of the early embryo. Cells grown in Serum/LIF, corresponding to the later pluripotent state, showed pronounced spreading on the surface and formed monolayers when cultured on a surface.

Closer inspection of the actin structure by STORM imaging showed characteristic differences in the nanoscale structure of the actin cytoskeleton between cells cultured in 2i and cells cultured in Serum/LIF. Cells cultured in Serum/LIF were observed to spread along the surface by flattening and extending out actin protrusions with dorsal stress fibers firmly connecting to the substrate. Stem cells cultured in 2i, on the other hand, mainly organize their actin in a cortical mesh with very few radial actin structures protruding from the cell and with less reinforcement along the cellular periphery than observed for cells cultured in Serum/LIF. The rounded shape of the individual cells cultured in 2i, as well as for the colonies they form, could be a result of low affinity for substrate attachment.

We probed the viscoelastic properties in the area close to the cells’ periphery, where we observed the significant geometrical differences in actin nano-architecture between the two culturing conditions. These results demonstrated a small difference at the cell periphery between the two culturing conditions, with the cells grown in Serum/LIF, which also displayed the most abundant actin structures along the cell periphery, being more elastic (stiffer) than cells cultured in 2i. It is reasonable that areas with more abundant actin polymers are more elastic, and this corresponds well with similar types of viscoelastic measurements done on *S. pombe* yeast cells, which also demonstrated a decrease in elasticity upon actin disruption (22).

The thick cortical actin in cells grown in 2i lying adjacent to the plasma membrane is likely to form tight cell junctions in 3D cell clusters by cell-cell adhesion through e-cadherin, thereby reinforcing the observed 3D dome-like structure. The question remains whether the actin structure could control the cell’s ability to differentiate by, e.g., regulation of the plasma membrane tension. The tension in membranes was recently found to regulate the fate of stem cells through tension regulation of endocytosis (37, 40). Our results show that stem cells already in early stages of differentiation begin to show distinct viscoelastic properties in the cell cortex, although these differences are still much smaller than at later stages in development (1). Stem cells grown in Serum/LIF media have abundant and dense actin fibers along the cellular periphery and more elastic material properties close to the cell periphery than the naïve cells, hence, there appears to be a stiffer cell cortex at this later developmental stage. It has been shown that inhibitor withdrawal from 2i media causes cells to decrease their membrane tension drastically as they transition towards differentiation (37, 40). While at first glance this might seem contrasting to our results, there are two important differences to notice: First, in this study we probe two separate stages of stem cell development; the slightly earlier and more naïve early blastocyst stage, recapitulated by culturing in 2i media, and the later primed or differentiated epiblast stage, recapitulated by cells cultured in Serum/LIF media (15). These are both defined and stable stages in our media conditions, whereas the studies reported in (37, 40) report on the transition from the naïve stage. Second, we focus on the actin structures located inside the membrane (not on the membrane itself), the presence of a more dense cortical actin network does not necessarily exclude lower membrane tension. As described in Bergert and others (40), during the transition the membrane tension decreases from a reduction in membrane-cytoskeleton linkage, and the transition could be inhibited by artificially linking the membrane to the cytoskeleton.

Here, we have shown that differences in the early developmental stages of ESCs are associated with dramatic differences in the actin nano-architecture of the cell cortex and also with changes in the physical properties both of the individual cells and of the colonies they form. Future studies will focus on implications of these differences on cell fate and on the mechanisms governing the interplay between the dynamic actin structure, the membrane, and cell fate regulation.

## Author Contributions

K.G.H. and Y.F.B built the super resolution imaging facility and performed all imaging experiments of the actin structure. I.I.P. and Y.F.B. performed the viscoelasticity measurements. Y.F.B. performed the data analysis. L.B.O. and P.M.B. designed and supervised the project. J.M.B. contributed with the cells and biological expertise. K.G.H. wrote the first draft and all authors commented and edited the manuscript.

## Acknowledgments

This work is financially supported by Danish National Research Foundation, grant DNRF116 (StemPhys), and the Danish Council for Independent Research, Natural Sciences, grant DFF-4181-00196.

## Supplementary Information

**Fig. S1:**
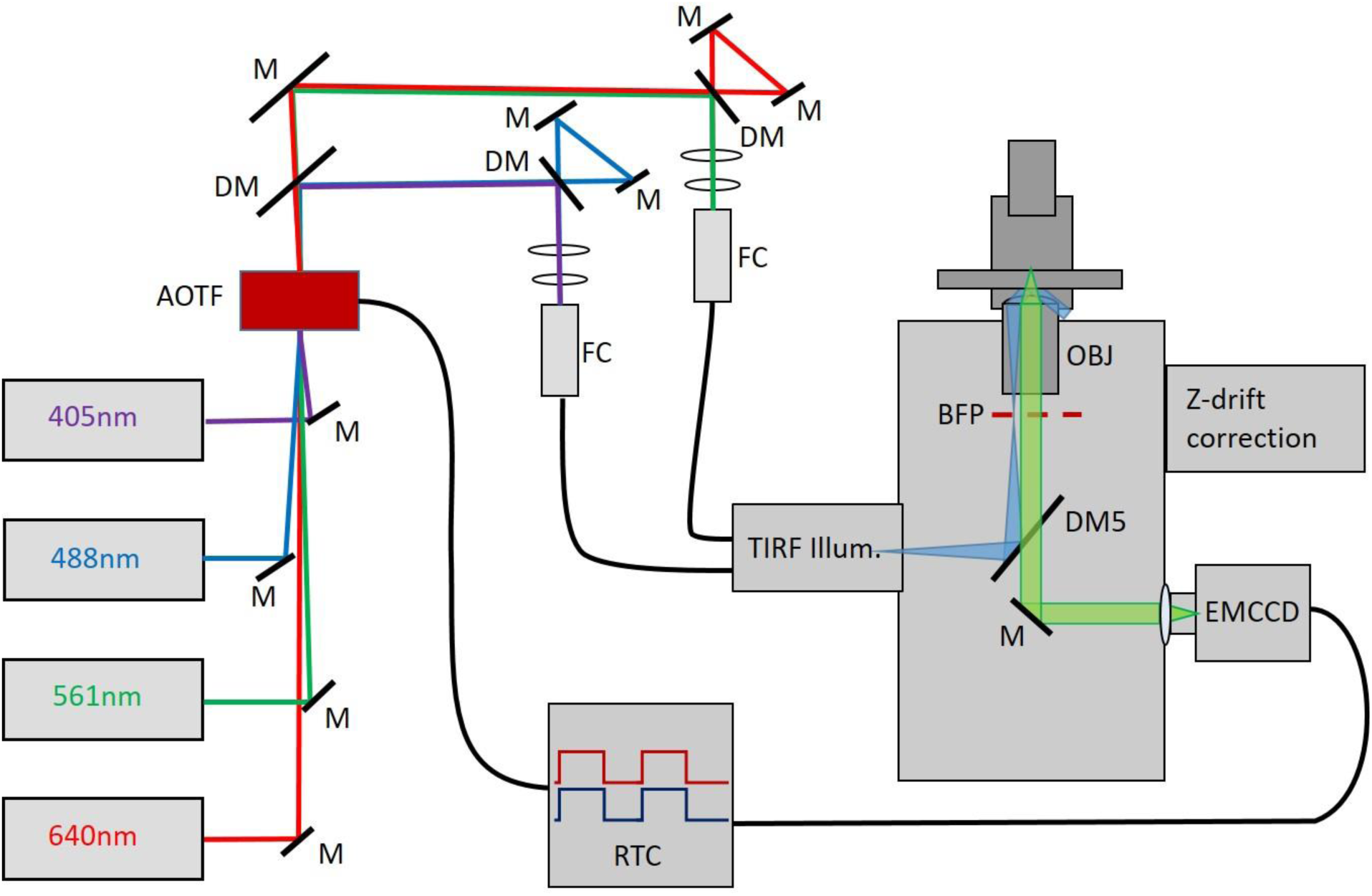
Sketch of the super-resolution setup. Lasers of wavelengths 405 nm (40 mW, Olympus), 488 nm (200 mW, Cobolt), 561 nm (200 mW, Cobolt), and 640 nm (200 mW, Toptica) are directed by mirrors through the aperture of an acousto-optic tunable filter (AOTF) (AA Opto Electronic). The beams are split by dichroic mirrors (DM) (AHF Analysentechnik), and the 405 and 488 nm beams are coupled into one single-mode optical fiber (FC) (Qioptic); likewise, the 561 and 640 nm beams are coupled into a second optical fiber. The TIRF illuminator allows motorized control of the focus position of the excitation light on the back focal plane (BFP) of the objective (OBJ) (1.45 NA, UAPON OTIRF, Olympus). Drift in the axial direction is continuously corrected by the Z-drift correction module (IX3-ZDC, Olympus). Images are captured on an EMCCD camera (ImagEM X2, Hamamatsu), which is coupled to the real-time controller (RTC, Olympus), that synchronizes the camera shutter with the AOTF in order to minimize sample illumination and bleaching.

**Fig. S2:**
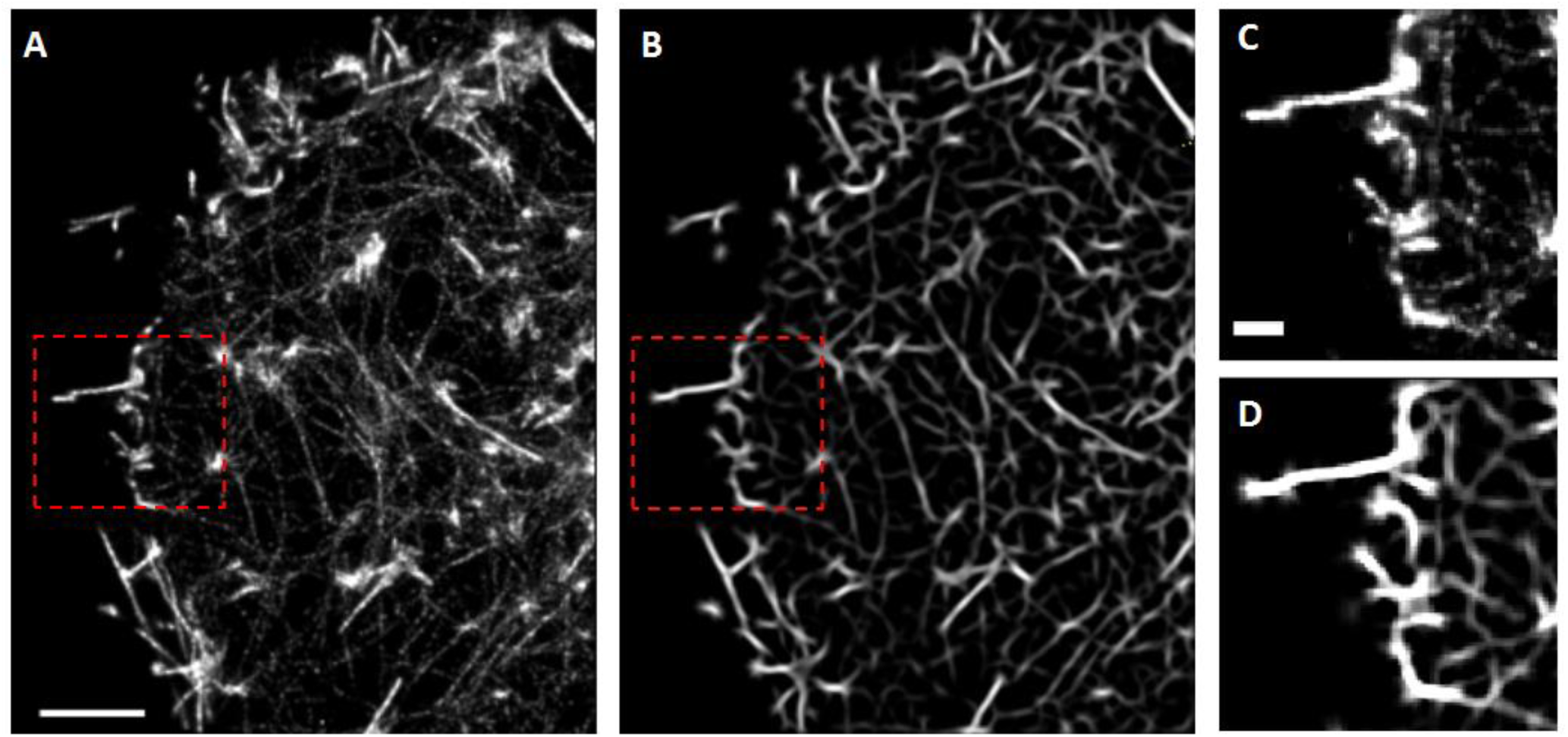
Spatial resolution enhancement**. (A)** Reconstructed STORM image of the actin cytoskeleton in a stem cell in 2i media. **(B)** Image corresponding to (A) after 2D Gabor filtering. (**C**-**D)** Magnified views of the red boxed regions in images (A) and (B), respectively. Scale bar shown in (A) is also valid in (B) and is 2 μm. Scale bar shown in (C) is also valid in (D) and is 0.5 μm.

## References

1. Urbanska, M., M. Winzi, K. Neumann, S. Abuhattum, P. Rosendahl, P. Müller, A. Taubenberger, K. Anastassiadis, and J. Guck. 2017. Single-cell mechanical phenotype is an intrinsic marker of reprogramming and differentiation along the mouse neural lineage. Development 144(23):4313–4321.

2. Engler, A. J., S. Sen, H. L. Sweeney, and D. E. Discher. 2006. Matrix Elasticity Directs Stem Cell Lineage Specification. Cell 126(4):677–689.

3. Wullkopf, L., A.-K. V. West, N. Leijnse, T. R. Cox, C. D. Madsen, L. B. Oddershede, and J. T. Erler. 2018. Cancer cells’ ability to mechanically adjust to extracellular matrix stiffness correlates with their invasive potential. Mol. Biol. Cell 29(20):2378–2385.

4. Han, Y. L., A. F. Pegoraro, H. Li, K. Li, Y. Yuan, G. Xu, Z. Gu, J. Sun, Y. Hao, S. K. Gupta, Y. Li, W. Tang, H. Kang, L. Teng, J. J. Fredberg, and M. Guo. 2019. Cell swelling, softening and invasion in a three-dimensional breast cancer model. Nature Physics.

5. Xia, S., Y. B. Lim, Z. Zhang, Y. Wang, S. Zhang, C. T. Lim, E. K. F. Yim, and P. Kanchanawong. 2019. Nanoscale Architecture of the Cortical Actin Cytoskeleton in Embryonic Stem Cells. Cell Reports 28(5):1251-1267.e1257.

6. Xu, K., H. P. Babcock, and X. Zhuang. 2012. Dual-objective STORM reveals three-dimensional filament organization in the actin cytoskeleton. Nat. Methods 9(2):185–188.

7. Han, B., R. Zhou, C. Xia, and X. Zhuang. 2017. Structural organization of the actin-spectrin–based membrane skeleton in dendrites and soma of neurons. Proceedings of the National Academy of Sciences 114(32):E6678.

8. Bongiorno, T., J. Gura, P. Talwar, D. Chambers, K. M. Young, D. Arafat, G. Wang, E. L. Jackson-Holmes, P. Qiu, T. C. McDevitt, and T. Sulchek. 2018. Biophysical subsets of embryonic stem cells display distinct phenotypic and morphological signatures. PloS one 13(3):e0192631.

9. Sim, Y. J., M. S. Kim, A. Nayfeh, Y. J. Yun, S. J. Kim, K. T. Park, C. H. Kim, and K. S. Kim. 2017. 2i Maintains a Naive Ground State in ESCs through Two Distinct Epigenetic Mechanisms. Stem cell reports 8(5):1312–1328.

10. Morgani, S. M., and J. M. Brickman. 2015. LIF supports primitive endoderm expansion during pre-implantation development. Development 142(20):3488.

11. Nichols, J., and A. Smith. 2009. Naive and Primed Pluripotent States. Cell Stem Cell 4(6):487–492.

12. Ying, Q.-L., J. Wray, J. Nichols, L. Batlle-Morera, B. Doble, J. Woodgett, P. Cohen, and A. Smith. 2008. The ground state of embryonic stem cell self-renewal. Nature 453(7194):519–523.

13. Martin Gonzalez, J., Sophie M. Morgani, Robert A. Bone, K. Bonderup, S. Abelchian, C. Brakebusch, and Joshua M. Brickman. 2016. Embryonic Stem Cell Culture Conditions Support Distinct States Associated with Different Developmental Stages and Potency. Stem Cell Reports 7(2):177–191.

14. Canham, M. A., A. A. Sharov, M. S. H. Ko, and J. M. Brickman. 2010. Functional heterogeneity of embryonic stem cells revealed through translational amplification of an early endodermal transcript. PLoS Biol. 8(5):e1000379–e1000379.

15. Morgani, S. M., M. A. Canham, J. Nichols, A. A. Sharov, R. P. Migueles, M. S. Ko, and J. M. Brickman. 2013. Totipotent embryonic stem cells arise in ground-state culture conditions. Cell Rep 3(6):1945–1957.

16. Betzig, E., G. H. Patterson, R. Sougrat, O. W. Lindwasser, S. Olenych, J. S. Bonifacino, M. W. Davidson, J. Lippincott-Schwartz, and H. F. Hess. 2006. Imaging intracellular fluorescent proteins at nanometer resolution. Science 313(5793):1642–1645.

17. Rust, M. J., M. Bates, and X. Zhuang. 2006. Sub-diffraction-limit imaging by stochastic optical reconstruction microscopy (STORM). Nat. Methods 3(10):793–795.

18. Heilemann, M., S. van de Linde, M. Schuttpelz, R. Kasper, B. Seefeldt, A. Mukherjee, P. Tinnefeld, and M. Sauer. 2008. Subdiffraction-resolution fluorescence imaging with conventional fluorescent probes. Angew Chem Int Ed Engl 47(33):6172–6176.

19. van de Linde, S., A. Loschberger, T. Klein, M. Heidbreder, S. Wolter, M. Heilemann, and M. Sauer. 2011. Direct stochastic optical reconstruction microscopy with standard fluorescent probes. Nat Protoc 6(7):991–1009.

20. Wullkopf, L., A. V. West, N. Leijnse, T. R. Cox, C. D. Madsen, L. B. Oddershede, and J. T. Erler. 2018. Cancer cells’ ability to mechanically adjust to extracellular matrix stiffness correlates with their invasive potential. Mol Biol Cell 29(20):2378–2385.

21. Andersen, T., A. Bahadori, D. Ott, A. Kyrsting, S. N. Reihani, and P. M. Bendix. 2014. Nanoscale phase behavior on flat and curved membranes. Nanotechnology 25(50):505101.

22. Tolić-Nørrelykke, I. M., E.-L. Munteanu, G. Thon, L. Oddershede, and K. Berg-Sørensen. 2004. Anomalous Diffusion in Living Yeast Cells. Physical Review Letters 93(7):078102.

23. Selhuber-Unkel, C., P. Yde, K. Berg-Sørensen, and L. B. Oddershede. 2009. Variety in intracellular diffusion during the cell cycle. Physical Biology 6(2):025015.

24. Hansen, P. M., I. M. Tolic-Nørrelykke, H. Flyvbjerg, and K. Berg-Sørensen. 2006. Tweezercalib 2.1.: Faster version of MatLab package for precise calibration of optical tweezers. Computer Physics Communications 175:572–573.

25. Wessel, Alok D., M. Gumalla, J. Grosshans, and Christoph F. Schmidt. 2015. The Mechanical Properties of Early *Drosophila* Embryos Measured by High-Speed Video Microrheology. Biophys. J. 108(8):1899–1907.

26. Norregaard, K., R. Metzler, C. M. Ritter, K. Berg-Sorensen, and L. B. Oddershede. 2017. Manipulation and Motion of Organelles and Single Molecules in Living Cells. Chemical reviews 117(5):4342–4375.

27. Ovesny, M., P. Krizek, J. Borkovec, Z. Svindrych, and G. M. Hagen. 2014. ThunderSTORM: a comprehensive ImageJ plug-in for PALM and STORM data analysis and super-resolution imaging. Bioinformatics 30(16):2389–2390.

28. Schindelin, J., I. Arganda-Carreras, E. Frise, V. Kaynig, M. Longair, T. Pietzsch, S. Preibisch, C. Rueden, S. Saalfeld, B. Schmid, J. Y. Tinevez, D. J. White, V. Hartenstein, K. Eliceiri, P. Tomancak, and A. Cardona. 2012. Fiji: an open-source platform for biological-image analysis. Nat. Methods 9(7):676–682.

29. Huang, B., W. Wang, M. Bates, and X. Zhuang. 2008. Three-dimensional super-resolution imaging by stochastic optical reconstruction microscopy. Science 319(5864):810–813.

30. Peishan Dai, Hanyuan Luo, Hanwei Sheng, Yali Zhao, Ling Li, Jing Wu, Yuqian Zhao, and K. Suzuki. 2015. A New Approach to Segment Both Main and Peripheral Retinal Vessels Based on Gray-Voting and Gaussian Mixture Model. PloS one 10:e0127748.

31. Hendargo, H. C., R. Estrada, S. J. Chiu, C. Tomasi, S. Farsiu, and J. A. Izatt. 2013. Automated nonrigid registration and mosaicing for robust imaging of distinct retinal capillary beds using speckle variance optical coherence tomography. Biomedical optics express 4(6):803–821.

32. Rezakhaniha, R., A. Agianniotis, J. T. C. Schrauwen, A. Griffa, D. Sage, C. V. C. Bouten, F. N. van de Vosse, M. Unser, and N. Stergiopulos. 2012. Experimental investigation of collagen waviness and orientation in the arterial adventitia using confocal laser scanning microscopy. Biomechanics and Modeling in Mechanobiology 11(3):461–473.

33. Püspöki, Z., M. Storath, D. Sage, and M. Unser. 2016. Transforms and Operators for Directional Bioimage Analysis: A Survey. Focus on Bio-Image Informatics. W. H. De Vos, S. Munck, and J.-P. Timmermans, editors. Springer International Publishing, Cham, pp. 69–93.

34. Inoue, S., V. Frank, M. Horning, S. Kaufmann, H. Y. Yoshikawa, J. P. Madsen, A. L. Lewis, S. P. Armes, and M. Tanaka. 2015. Live cell tracking of symmetry break in actin cytoskeleton triggered by abrupt changes in micromechanical environments. Biomaterials science 3(12):1539–1544.

35. Gupta, M., B. R. Sarangi, J. Deschamps, Y. Nematbakhsh, A. Callan-Jones, F. Margadant, R.-M. Mège, C. T. Lim, R. Voituriez, and B. Ladoux. 2015. Adaptive rheology and ordering of cell cytoskeleton govern matrix rigidity sensing. Nature Communications 6(1):7525.

36. Chalut, Kevin J., and Ewa K. Paluch. 2016. The Actin Cortex: A Bridge between Cell Shape and Function. Dev. Cell 38(6):571–573.

37. De Belly, H., P. H. Jones, E. K. Paluch, and K. J. Chalut. 2019. Membrane tension mediated mechanotransduction drives fate choice in embryonic stem cells. bioRxiv:798959.

38. Tojkander, S., G. Gateva, and P. Lappalainen. 2012. Actin stress fibers – assembly, dynamics and biological roles. J. Cell Sci. 125(8):1855–1864.

39. Naumanen, P., P. Lappalainen, and P. Hotulainen. 2008. Mechanisms of actin stress fibre assembly. Journal of Microscopy 231(3):446–454.

40. Bergert, M., S. Lembo, D. Milovanović, M. Börmel, P. Neveu, and A. Diz-Muñoz. 2019. Cell surface mechanics gate stem cell differentiation. bioRxiv:798918.

